# Biolistics-mediated transformation of hornworts and its application to study pyrenoid protein localization

**DOI:** 10.1101/2023.10.23.563637

**Authors:** Declan J. Lafferty, Tanner A. Robison, Andika Gunadi, Laura H. Gunn, Joyce Van Eck, Fay-Wei Li

## Abstract

Hornworts are a deeply diverged lineage of bryophytes that are sister to mosses and liverworts. Hornworts have an array of unique features that can be leveraged to illuminate not only the early evolution of land plants, but also alternative paths for nitrogen and carbon assimilation via cyanobacterial symbiosis and a pyrenoid-based CO_2_-concentrating mechanism (CCM), respectively. Despite this, hornworts are one of the few plant lineages with limited available genetic tools. Here we report an efficient biolistics method for generating transient-expression and stable transgenic lines in the model hornwort, *Anthoceros agrestis*. An average of 569 (± 268) cells showed transient expression per bombardment, with green fluorescent protein expression observed within 48 hours. A total of 81 stably transformed lines were recovered across three separate experiments, averaging six lines per bombardment. We followed the same method to transiently transform nine additional hornwort species, and obtained stable transformants from one. This method was further used to verify the localization of Rubisco and Rubisco activase in pyrenoids, which are central proteins for CCM function. Together, our biolistics approach offers key advantages over existing methods as it enables rapid transient expression and can be applied to widely diverse hornwort species.

**Highlight:** We developed a new transformation method for hornworts, a lineage of understudied bryophytes, and demonstrated its effectiveness in studying subcellular localization of proteins involved in pyrenoid-based CO_2_-concentrating mechanism.

## Introduction

Hornworts are a unique group of bryophytes that exhibit distinct traits among land plants. Evolutionarily, hornworts diverged from the closest extant clade, setaphytes (mosses and liverworts), over 480 million years ago (Morris et al., 2018). Many of hornworts’ developmental, anatomical, and physiological features are not found in other plant lineages. As such, hornworts are of paramount importance to retrace plant evolutionary history (Li et al., 2020; Frangedakis et al., 2021a; Frangedakis et al., 2021b).

Hornworts have evolved unconventional ways to source nitrogen and carbon. For nitrogen, all hornwort species are known to engage in a symbiotic relationship with nitrogen-fixing cyanobacteria (Meeks, 1998; Nelson et al., 2021). Symbiotic cyanobacteria are hosted inside specialized slime cavities in the hornwort thallus, where they differentiate into heterocysts, a cell type dedicated to nitrogen fixation. While several hornwort genes have been recently identified that might be involved in symbiosis regulation or maintenance (Li et al., 2020; Chatterjee et al., 2022), they have not been functionally characterized.

For carbon capture, hornworts are the only land plants that evolved a pyrenoid-based CO_2_-concentrating mechanism (CCM). In this case, the CO_2_-fixing enzyme, ribulose-1,5-bisphosphate carboxylase/oxygenase (Rubisco), is compartmentalized inside pyrenoids, which are special chloroplast structures sandwiched by thylakoids (Vaughn et al., 1992). Through the coordinated actions of bicarbonate transporters and carbonic anhydrases, CO_2_ is concentrated within pyrenoids thereby boosting the efficiency of Rubisco (Li et al., 2017; He et al., 2023). Interestingly, pyrenoids (and the associated CCM) have been repeatedly gained and lost during hornwort evolution (Villarreal & Renner, 2012), providing unique evolutionary replications for examining the underlying genetic mechanisms. It has been hypothesized that engineering a biophysical CCM into crop plants could enhance photosynthetic efficiency (Adler et al., 2022), but attempts to transfer algal pyrenoid-based CCM are challenged by the incompatibility between land plant Rubisco and algal pyrenoid components (Meyer et al., 2016; Atkinson et al., 2020). On the other hand, hornwort Rubisco and chloroplasts are both evolutionarily and structurally similar to those in crop plants, providing a CCM that may be more readily engineered into crops. Currently, none of the genetic components of hornwort pyrenoid structure and development have been identified, outside of Rubisco.

*Anthoceros agrestis* has recently emerged as a potential model hornwort species, with a sequenced genome and an optimized tissue culture growth medium (Szövényi et al., 2015; Li et al., 2020; Gunadi et al., 2022). Its life cycle is dominated by a haploid gametophyte stage, making it an ideal target for genetic engineering. However, the lack of transformation tools has hampered the progress of functional genomics in hornworts. Recently, an *Agrobacterium-*mediated transformation method was reported for *A. agrestis* and has been successfully used for a number of hornwort species (Frangedakis et al., 2021b; Waller et al., 2023). Transient transformation of hornwort protoplasts has also been reported (Neubauer et al., 2022), although thalli regeneration from protoplasts has not been possible. The biolistics method, also known as particle bombardment or gene gun technology, is another powerful tool used for genetic engineering (Sanford et al., 1987) and that has not been explored for hornworts. This method involves the introduction of microscopic particles coated with DNA into plant cells.

Biolistics has a number of advantages over other transformation methods. First, it is relatively simple and requires less technical expertise. The preparation time for biolistics is relatively fast, taking less than a couple of hours for particle preparation and subsequent transformation. *Agrobacterium-*mediated methods involve the use of live bacteria and co-cultivation steps, and protoplast transformation requires cell wall digestion, all of which can be difficult to optimize (Lacroix & Citovsky, 2020). Second, biolistics can deliver a wider range of cargoes into plant cells including plasmid DNA, linearized DNA (such as PCR products), RNA, ribonucleoproteins, and proteins (Fu et al., 2000; Martin-Ortigosa & Wang., 2014; Svitashev et al., 2016; Zhang et al., 2016). Finally, biolistics can be used to transform plastids (Boynton et al., 1988; Svab et al., 1990), which is crucial to study the genes involved in pyrenoid function. Conversely, the downsides of the biolistics method include the possibility to cause severe cellular damage and the method may require expensive supplies and equipment. Reports have also shown a higher number of integrated gene copies when compared with *Agrobacterium*-mediated transformation, potentially causing genetic instability and gene silencing (Kohli et al., 2003). The extent of these impacts, however, has not yet been explored in hornworts.

In this study, we describe the development of a biolistics transformation method for hornworts, demonstrating both transient and stable transformation of *A. agrestis* and *A. fusiformis*, as well as transient transformation of nine other species across the hornwort tree of life. These transformable species share the most recent common ancestor almost 300 million years ago, before the origin of flowering plants. Using our optimized method, we were able to observe transient expression of the green fluorescent protein (GFP) reporter gene within two days of transformation, and recover stable transgenic lines after 8-10 weeks. Importantly, we applied this method to examine the subcellular localization of Rubisco and Rubisco activase, both of which are key elements in CCMs, which represents the first gene functional study in hornworts, and allows for comparative research across bryophytes and vascular plants.

## Materials and Methods

### Plant tissue preparation

*Anthoceros agrestis* cultures were grown axenically on AG medium supplemented with 0.2% sucrose at 22°C under a 16/8 h light/dark cycle with 6-25 µmol/m²/s light intensity provided by Ecolux XL Starcoat F32T8 XL SP30 ECO paired with F32T8 XL SP41 ECO fluorescence bulbs (General Electric, USA) as described in Gunadi et al. (2022). Thallus tissue (haploid) was prepared for transformation by placing 3-6 g (fresh weight) of tissue in a 50 mL Falcon tube containing 30 mL of sterile deionized water, containing 30 µL of 300 mg/mL timentin (final concentration of 300 mg/L) (T-104-25, Goldbio, USA), followed by homogenization with a T18 digital ULTRA TURRAX homogenizer (IKA, Germany) for 5 s at 4000 rpm. The tissue was filtered using a 100 µM cell strainer (C4100, MTC Bio, USA) and washed with 50 mL sterile deionized water. Approximately 200 mg of tissue was plated on AG medium, supplemented with 2% sucrose. Tissue was spread as thinly as possible in a 2.5 cm circle using a Nunc cell scraper (ThermoFisher Scientific, USA). The plates were left open for 10 min to remove excess moisture. Tissue was allowed to recover for 7 days before transformation. These procedures were done in a laminar flow hood.

### DNA construct preparation

Constructs were designed using the OpenPlant toolkit (Sauret-Güeto et al., 2020). The Aa049 construct was assembled using the following L0 parts: Native *A. agrestis Elongation factor 1 alpha* (*Ef1α*) promoter, OP_049 35S promoter, OP-020 *hygromycin phosphotransferase* (*hpt)*, OP-023 *eGFP*, OP_037 *CTAG_Lti6b*, OP-053 *NoS* terminator. Vector ligations and *E. coli* transformation were performed as described in Frangedakis et al. (2021b). To obtain high concentrations of plasmid DNA, 100 ml of *E. coli* culture was grown overnight at 37°C. The plasmid DNA was extracted using the PureYield™ Plasmid Midiprep System (A2492, Promega, USA), followed by vacuum concentration in a SpeedVac Concentrator (Thermofisher Scientific, USA) to obtain DNA concentrations of 1 µg/µL.

### Gold particle preparation

To prepare for 10 separate bombardments, a total of 20 µL of 1 µg/µL plasmid DNA was combined with 50 µL of gold microparticles (1 µm, 50 mg/ml, suspended in 50% (v/v) sterile glycerol) (#1652263, Bio-Rad, USA), 100 µL 2.5 M CaCl_2_ and 40 µL 0.1 M spermidine (S0266, Sigma-Aldrich, USA), under constant vortexing. The DNA/gold mixture was vortexed for 1 min before being pelleted by centrifugation at 3500 x g for 8 s. The supernatant was removed and 180 µL of 70% (v/v) ethanol was added and vortexed for 1 min. The DNA/gold mixture was centrifuged again at 3500 x g for 8 s. The supernatant was removed and 200 µL of 100% ethanol were added and vortexed for 1 min, followed by another centrifugation step and the removal of 135 µL of supernatant. The DNA/gold mixture was resuspended in a Branson 2510 Ultrasonic Cleaner and 5 µL was spread onto each sterilized macrocarrier (#1652335, Bio-Rad, USA) and left to dry for 30 min.

### Biolistics transformation

Biolistics was performed using the Particle Delivery System PDS-1000/He (#1652257, Bio-Rad, USA). Thallus tissue was bombarded with a target distance of 14 cm, under a vacuum of 28 inHg. The burst pressure was generated using a 450 psi rupture disk (#1652326, Bio-Rad, USA). The macrocarrier and rupture disc were retained using a stopping screen (#1652257, Bio-Rad, USA). The tissue was left to recover for 7 days.

### Imaging and quantification of transient transformation efficiency

Transient transformation efficiency was estimated using a pipeline developed in CellProfiler (v4.2.5) (Stirling et al., 2021). Fluorescent images of transformed tissues were taken 3 days post bombardment, using a GPF filter on a Leica M205 stereomicroscope at a magnitude of 10 mm x 7.5 mm. Nine images were taken to encompass all tissue used for transformation. The exposure time was set between 10 – 12 s, depending on transient GFP fluorescent intensity, and was kept constant for all images within an experiment. The default pipeline parameters were used with adjustments made to cell diameter (between 6- and 30-pixel units in diameter) and object intensity (green intensity between 0.1 and 0.9). Red signal represents cells expressing GFP and blue signals are objects classified by CellProfiler that are not expressing GFP; these may be artifacts or untransformed cells/tissue. The pipeline can be found on Figshare (10.6084/m9.figshare.24243559). The number of transients in each photo was combined to show the total transients in each replicate.

### Regeneration of stably transformed lines

To regenerate stably transformed lines, tissue from all replicates was pooled and homogenized (as described above) 7 days post bombardment. The homogenized tissues were collected using a 100 µM cell strainer, washed with sterile water, and thinly spread on 70 mm diameter Whatman filter paper (WHA1001070, Sigma-Aldrich, USA) overlain on solid AG medium supplemented with 0.2% sucrose. The number of plates prepared corresponds to the number of replicates pooled. This tissue was left to recover for 3 days before transfer to AG medium supplemented with 0.2% sucrose and 10 mg/L hygromycin (#10687010; Invitrogen, USA), by moving the filter paper containing the tissue. After 4 weeks the tissue was transferred again to fresh AG medium supplemented with 0.2% sucrose and 10 mg/L hygromycin. Stably transformed thallus tissue is visible within 8-10 weeks of transformation. Images were taken on a Leica M205 stereomicroscope and a Leica TCS SP5 Laser Scanning Confocal Microscope.

**Figure 1.**
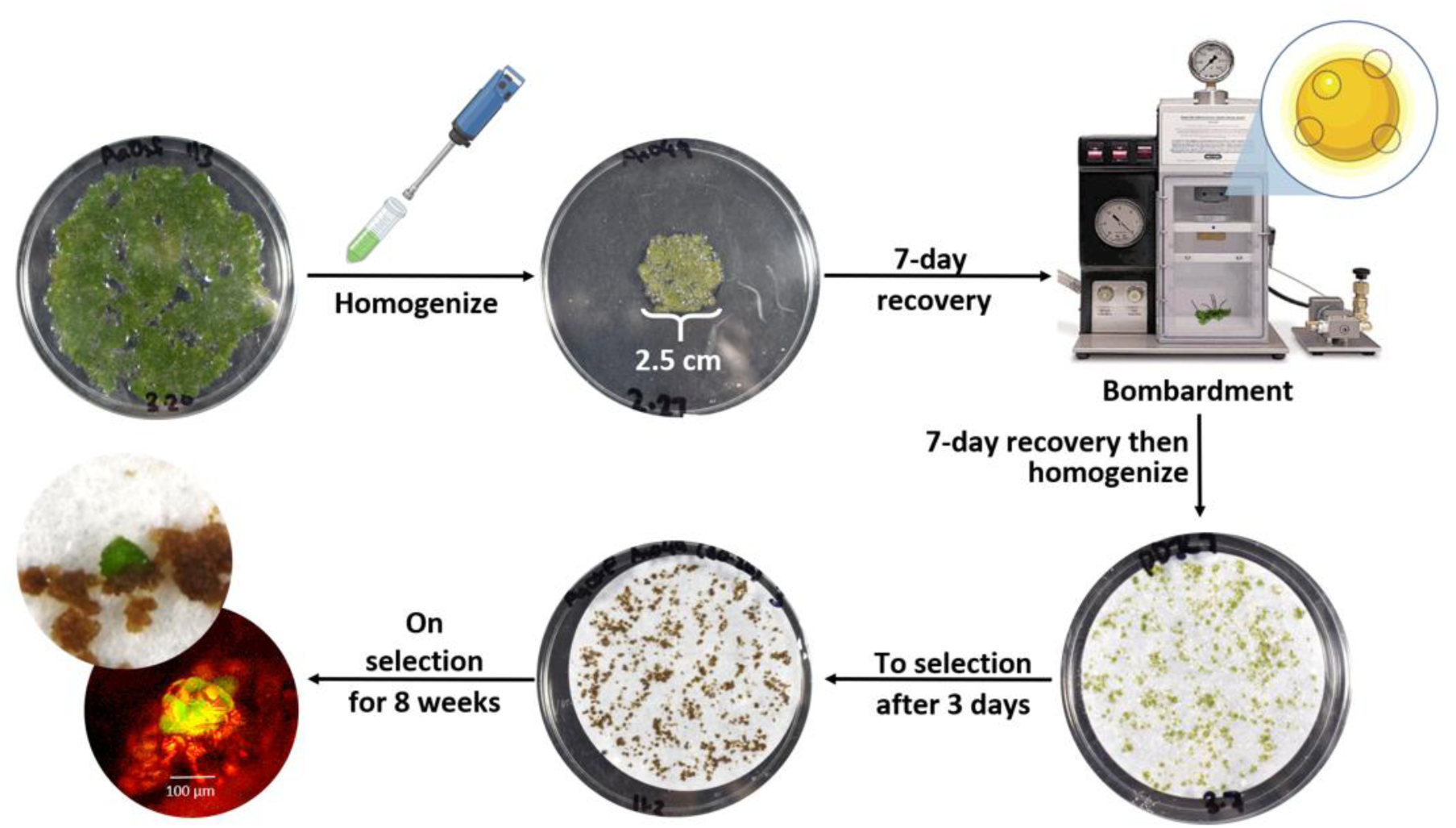
Biolistics method overview. Healthy *A. agrestis* thallus tissue, grown on AG medium containing 0.2% sucrose, is collected and homogenized. Tissue is washed, filtered, and placed in a 2.5 cm diameter circle onto AG medium containing 2% sucrose. The tissue was allowed to recover for 7 days both before and after transformation. The tissue is pooled from all replicates, homogenized again and placed onto filter paper on AG medium containing 0.2% sucrose to recover for 3 days. For antibiotics selection, tissue is transferred to AG medium, containing 0. 2% sucrose and 10 mg/L hygromycin. Transformed thallus tissue is visible approximately 8-10 weeks after transformation.

### Genome sequencing and transgene copy number estimation

A total of 12 stable lines (four per experiment) were shotgun sequenced to infer transgene copy number. *A. agrestis* thallus tissue was flash-frozen with liquid nitrogen and ground to a fine powder using a mortar and pestle. DNA was extracted using the CTAB protocol described in Li et al. (2020). Illumina library preparation and paired-end sequencing (150 bp), with 50X coverage, was performed by Novogene Corp. Read trimming and quality check were done using fastp v0.23.3 (Chen et al., 2018). We estimated transgene copy number by calculating the median read depth ratio between the *35S* promoter:GFP sequence and single-copy genes in the genome. Using Benchmarking Universal Single-Copy Orthologs (BUSCO; v5.4.7) (Manni et al., 2021), we selected 375 single-copy genes (Supp. Table 1) from *A. agrestis* for read-depth comparison. BWA (v0.7.17) (Li & Durbin, 2009) and SAMtools (v1.17) (Danecek et al., 2021) were used to map reads and calculate read depth, respectively, for the BUSCO genes and the transgene. The command lines and program parameters can be found on figshare (10.6084/m9.figshare.24243559).

**Table 1.** Transgene copy number estimate for the sequenced stable lines. Note that although two lines had a BUSCO-GFP ratio of less than 1, GFP expression was still observed and there was even read coverage across the entire GFP cassette with no missing sequences.

|  | Line | BUSCO<br>Median Read<br>Depth | GFP Cassette<br>Median Read<br>Depth | BUSCO-GFP<br>Ratio | Inferred copy<br>number |
| --- | --- | --- | --- | --- | --- |
| Experiment 1 | 1a | 53 | 370 | 6.98 | 7 |
|  | 1b | 41 | 552 | 13.46 | 13 |
|  | 1c | 53 | 1552 | 29.28 | 29 |
|  | 1d | 42 | 23 | 0.55 | 1 |
| Experiment 2 | 2a | 50 | 50 | 1.00 | 1 |
|  | 2b | 67 | 133 | 1.99 | 2 |
|  | 2c | 58 | 45 | 0.78 | 1 |
|  | 2d | 74 | 5541 | 74.88 | 75 |
| Experiment 3 | 3a | 48 | 153 | 3.19 | 3 |
|  | 3b | 44 | 45 | 1.02 | 1 |
|  | 3c | 28 | 1826 | 65.21 | 65 |
|  | 3d | 42 | 422 | 10.05 | 10 |

### Transformation of additional hornwort species

To test the applicability of our method in other hornworts, we selected nine species spanning seven genera and all of the five hornwort families (*Anthoceros fusiformis, Anthoceros punctatus, Anthoceros tuberculatus, Leiosporoceros dussii, Megaceros flagellaris, Notothylas orbicularis, Phaeoceros* sp*., Phaeomegaceros chiloensis* and *Phymatoceros phymatodes*). Cultures were maintained as described earlier. Transformation of these species followed the same method used for *A. agrestis*, using the Aa049 construct.

### Fluorescent protein-tagging and transformation

The *Rubisco small subunit* (*RbcS*, AagrOXF_evm.model.utg000008l.255.1) and *Rubisco activase* (*RCA*, AagrOXF_evm.model.utg000011l.50.1) CDS were identified in the *A. agrestis* genome (Li et al., 2020) using Orthofinder (v2.5.4) (Emms & Kelly, 2019). In the case of *RbcS*, multiple copies of the gene were present, so a copy that showed consistently high expression across multiple conditions was selected to ensure that it was a functional isoform (Li et al., 2020). DNA sequences were synthesized by Twist Biosciences and cloned into L1 vectors. *RCA*, tagged with the *mVenus* fluorescent protein, which was under the control of the *EF1*α and had a *NoS* terminator. *RbcS,* tagged with *mScarlet-I,* was under the control of the *A. agrestis RbcS* promoter and also had a *NoS* terminator. The L1 construct was combined with the *Ef1α:hpt:NoS* construct in the *pCSA* L2 vector. Plasmid maps and sequences can be found on figshare (10.6084/m9.figshare.24243559). A*. agrestis* transient and stable transformation followed the method described earlier.

## Results

### Efficient transient transformation by biolistics

After conducting experiments to refine the biolistics method for *A. agrestis*, it was determined that the optimal method parameters encompassed the use of 1 µm gold particles, a helium pressure of 600 psi, a vacuum pressure of 28 inHg, and a bombardment plate distance of 14 cm from the source. These parameters were chosen for future experiments because more severe tissue damage was observed when bombarding at higher pressures and shorter plate distances. To quantify the number of cells transiently expressing GFP, we developed an efficient semi-automatic procedure using CellProfiler (Stirling et al., 2021). This method has previously been used to quantify the transient transformation efficiency in biolistically transformed onion and *Nicotiana benthamiana* (Miller et al., 2021). We confirmed the results from CellProfiler with manual counting and found a strong linear correlation, with a Pearson correlation coefficient (r) of 0.98 (Supp. Fig. 1).

To assess the efficacy and reproducibility of transient transformation, five separate experiments (with 10 replicates each) were performed (Fig. 2C). Across all successfully transformed replicates, there was an average of 569 (± 268) transiently expressing GFP cells from one bombardment experiment. These results were quantified 72 hours after transformation, with fluorescence first observed within 24 hours. We found high variability between replicates in each experiment, for example in experiment 1, one replicate had 1158 transiently expressing cells while another had 114. This variability was reduced when comparing the means from each experiment, which ranged from 474 to 701 transient events (Fig. 2). In addition, not all replicates were successfully transformed (i.e. no GFP signal observed from any cell). These were removed from downstream analysis (2-4 per experiment; Fig. 2). It is unknown what caused inconsistencies, with potential factors being poor tissue health, tissue moisture level or aggregation of gold particles on the macrocarrier, all of which can reduce transformation efficiency (Southgate et al., 1995).

**Figure 2.**
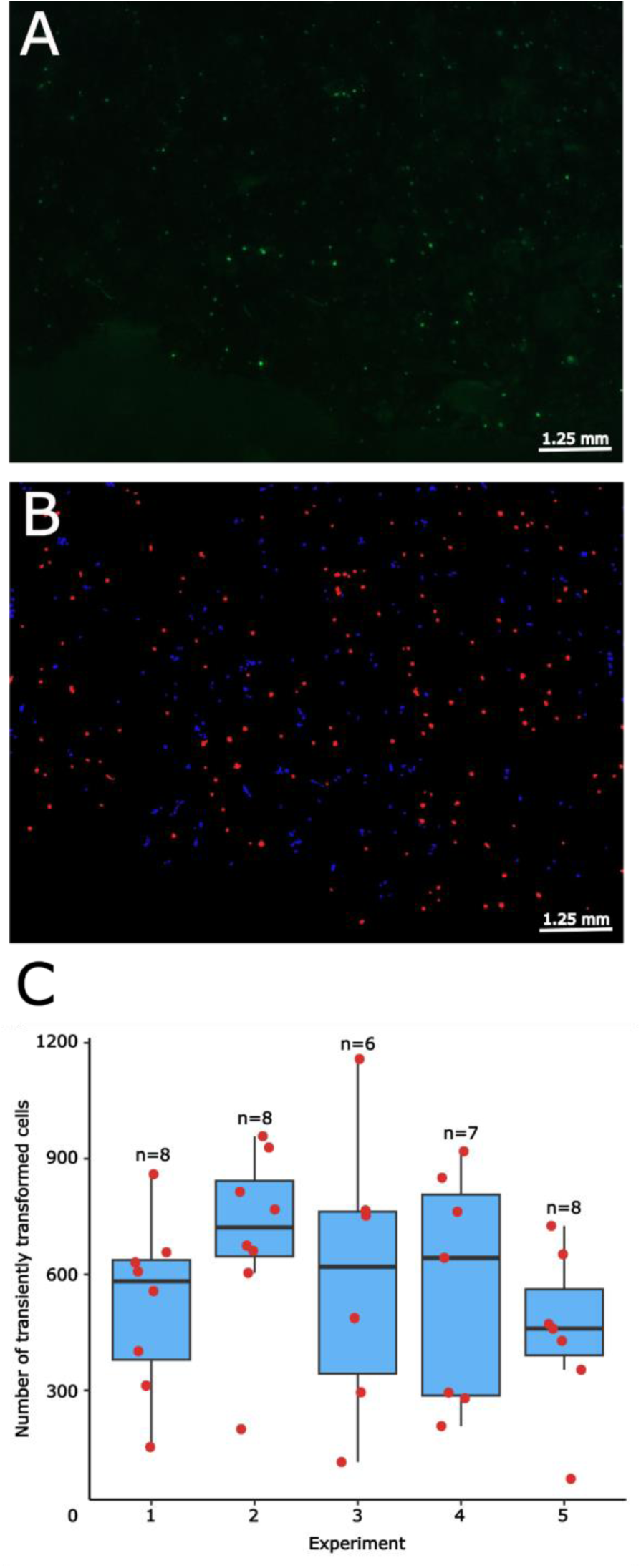
Transient transformation of *Anthoceros agrestis* and event quantification. (A) The unprocessed image of tissue transiently expressing GFP, taken with a Leica M205 stereomicroscope. (B) The same image in A after processing with CellProfiler, where blue signals represent artifact objects or cells/tissue not expressing GFP and red signals are transformed cells expressing GFP. (C) Boxplot displaying the number of cells transiently expressing GFP, across five separate experiments. Individual replicates are plotted as red dots. The box represents the 25th and 75th percentile, with the horizontal line being the median value and the vertical lines marking the minimum and maximum, excluding outliers.

### Recovery of stable transformants

Although transient transformation resulted in hundreds of transient events, it was initially challenging to recover stably transformed lines. In our earlier attempts, we allowed the bombarded tissue to recover for 7 days before cutting into 2 mm^2^ pieces and transferring to AG medium containing 0.2% sucrose and 10 mg/L hygromycin. Over the following weeks, we observed that GFP-expressing tissue persisted, however, the surrounding tissue also survived much longer than other non-transformed tissue. HPT inactivates hygromycin via phosphorylation (Rao et al., 1983) and in *Arabidopsis*, HPT was shown to be secreted into the extracellular space (Zhang et al., 2011). It is possible that hygromycin is inactivated in the medium surrounding transformed tissue, allowing untransformed tissue to ‘feed’ off it, limiting regeneration. To overcome this, tissue was homogenized before being placed on selection medium in order to reduce the amount of non-transformed tissue surrounding the transformed tissue. After 8-10 weeks of selection, the first regenerating hygromycin-resistant thallus tissue was visible and were transferred to fresh selection medium for propagation. The presence of the transgene was confirmed via fluorescent microscopy and PCR, with primers designed to span the *Ef1α* promoter and *hpt* CDS (Supp. Fig. 2-4). This was repeated across three separate experiments, with five replicates in experiments 1 and 2, and four replicates in experiment 3. These experiments regenerated 20, 49 and 12 individual lines, respectively, totaling 81 lines. On average, each bombardment of 0.2 g of tissue yielded six stable transformants.

**Figure 3.**
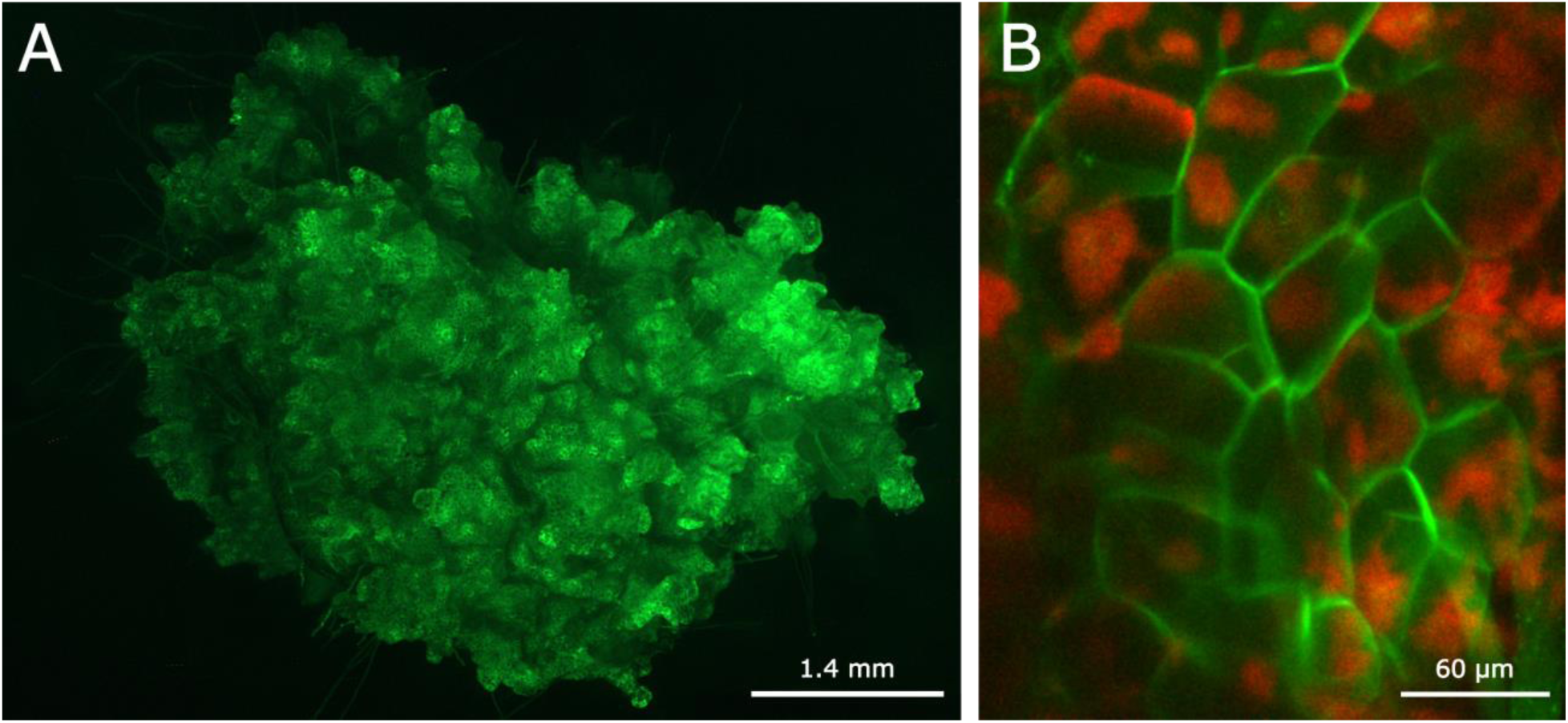
Stable transformation of *Anthoceros agrestis*. Transformed thallus expressing membrane-localized GFP was imaged using (A) a Leica M205 stereomicroscope, with the GFP channel, and (B) a Leica TCS SP5 Laser Scanning Confocal Microscope, showing both chlorophyll autofluorescence in red and GFP signal in green.

A drawback of biolistics-mediated transformation is the potential for high transgene copy number, which may lead to instability and potential gene silencing (Kohli et al., 2003). To assess the transgene copy number, we shotgun sequenced four stable lines per experiment, and compared the median read depth of the *35Spro:GFP* sequence to the median read depth for a set of single-copy genes (here the BUSCO genes were used; Table 1, Supp. Table 1). The whole plasmid was not used for read depth comparison as it contained the native hornwort Ef1a promoter and multiple copies of the NOS terminator. We found that 50% of the sequenced lines had a copy number fewer than four, and 30% had a single copy. Several lines had very large copy numbers, with the highest estimated at 75.

### Broad applicability to other hornwort species

To assess the applicability of the biolistics method to other hornwort species, we tested a total of nine species spanning the entire hornwort phylogeny: *Anthoceros fusiformis, Anthoceros punctatus, Anthoceros tuberculatus, Leiosporoceros dussii, Megaceros flagellaris, Notothylas orbicularis, Phaeoceros* sp*., Phaeomegaceros chiloensis* and *Phymatoceros phymatodes*. The majority of species had fewer transiently expressing GFP cells than *A. agrestis*, while both *A. punctatus* and *N. orbicularis* had a greater number (Fig. 4). Stable transformants were obtained for *A. fusiformis* with 29 lines recovered from one experiment (Supplementary Fig. 5). The method will need to be optimized for each species in order to increase the efficiency of transient transformation and to obtain stable transformants.

**Figure 4.**
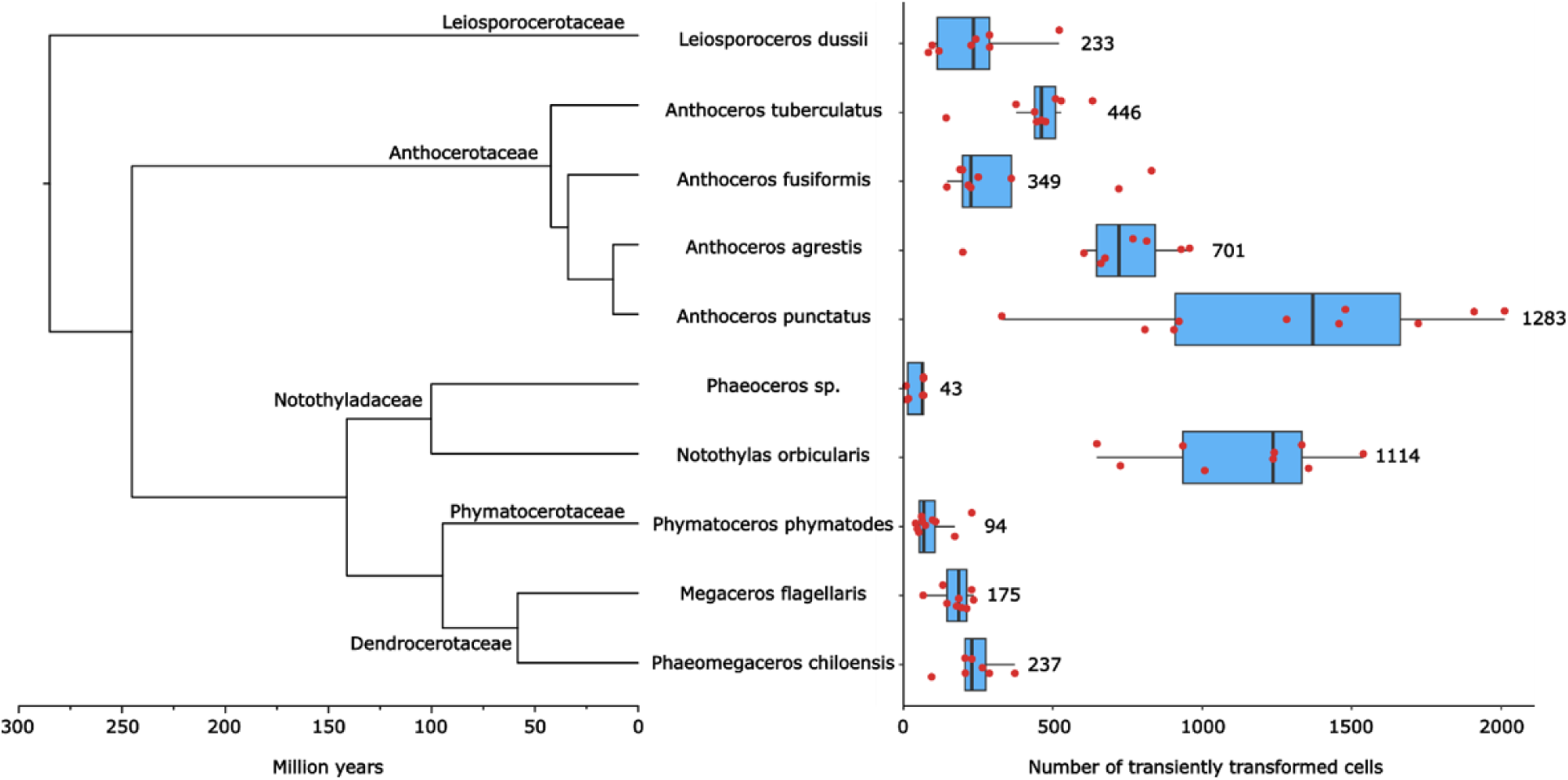
Transient transformation of hornwort species across the phylogeny. Our method can be directly applied to generate transient transformants in species covering all hornwort families and most of the genera. Phylogenetic tree displays timing of divergence, adapted from Bechteler et al. (2023). Red dots represent the value of replicates and numbers represent the average number of transiently transformed cells per bombardment. The box represents the 25th and 75th percentile, with the vertical line being the median value. The horizontal lines represent the minimum and maximum, excluding outliers.

### Transient and stable expression of fluorescently tagged Rubisco and Rubisco activase

To demonstrate the applicability of the biolistics-mediated transformation method for hornwort studies, we used it to investigate the structure of hornwort pyrenoids by localizing two putative pyrenoid-containing proteins, the Rubisco small subunit (RbcS) and Rubisco activase (RCA) through co-transformation of *A. agrestis*. The *RbcS* coding sequence was fused to the mScarlet-I fluorophore and the *RCA* coding sequence was fused to the mVenus fluorophore. Co-transformation of these two constructs clearly showed that RbcS and RCA co-localized, with fluorescence tightly confined within round structures in the chloroplast corresponding to hornwort pyrenoids (Fig. 5). Pyrenoids appear as distinct black holes under the chlorophyll autofluorescence channel, which is consistent with the early ultrastructural studies based on transmission electron microscopy (Vaughn et al., 1992). Furthermore, we obtained stably transformed lines expressing either the RbcS:msScarlet-I or the RCA:mVenus constructs. Again, the fluorescence signals were highly concentrated in pyrenoid structures that are devoid of chlorophyll autofluorescence.

**Figure 5.**
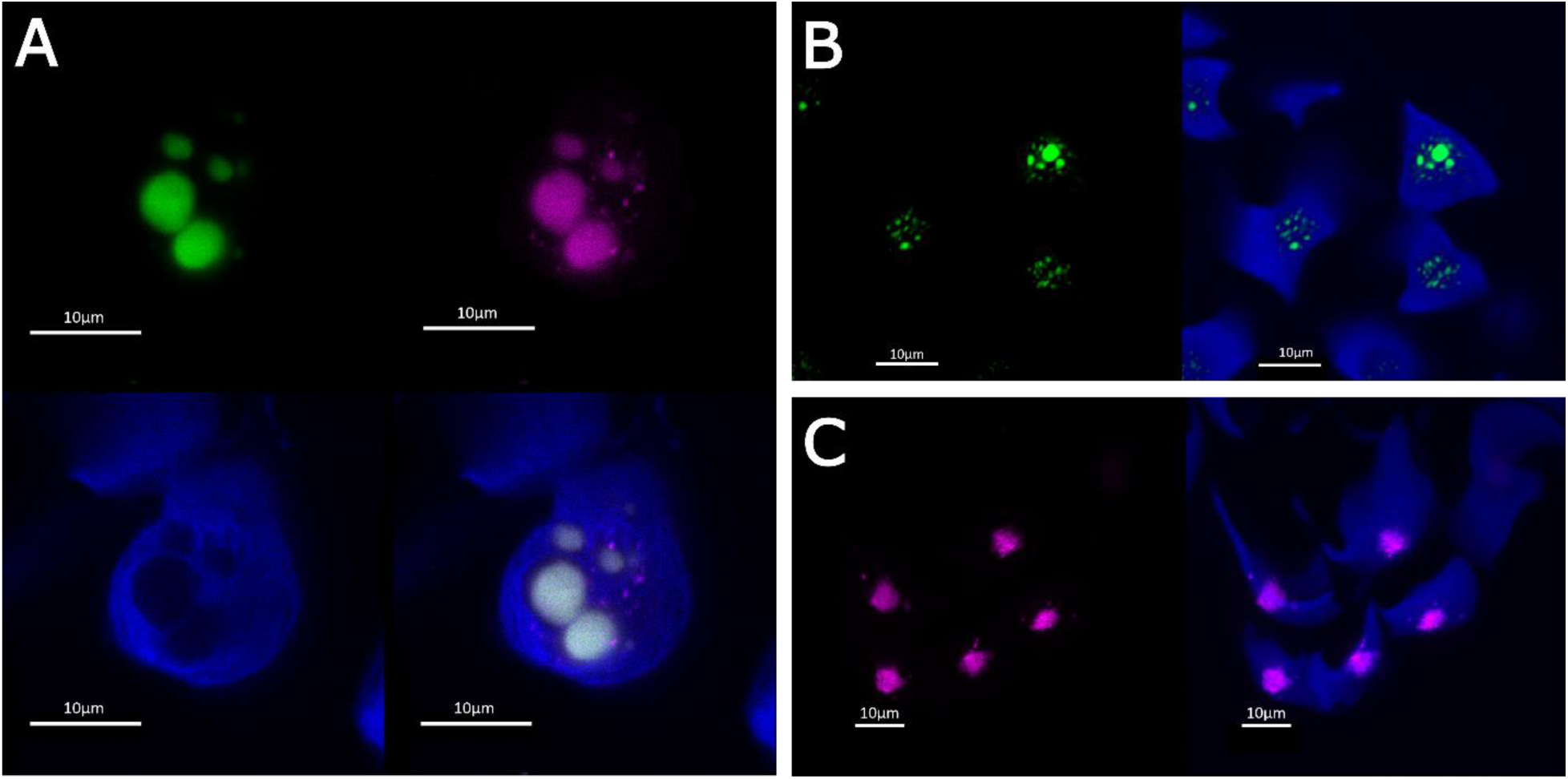
Transient and stable transformation of pyrenoid-related genes in *Anthoceros agrestis*. (A) *A. agrestis* thallus tissue was co-transformed with *RbcS:mScarlet-I* and *RCA:mVenus* constructs. The pyrenoid regions lack chlorophyll autofluorescence (blue) and are instead packed with tagged RbcS and RCA. The overlapping mScarlet-I and mVenus fluorescence signal strongly suggests co-localization of RbcS and RCA to pyrenoids. *A. agrestis* thallus tissue stably expressing (B) *RCA:mVenus* and (C) *RbcS:mScarlet-I*, with chlorophyll in blue. All images were taken using a Leica TCS SP5 Laser Scanning Confocal Microscope.

## Discussion

### A biolistics method for transient and stable transformation across the hornwort phylogeny

Our current method resulted in a high number of transient and stable events for *A. agrestis*. From 0.2 g of bombarded tissue, we obtained on average 569 (± 268) transiently transformed cells and six stable insertion lines. Furthermore, we provide the first account of transient transformation in *A. tuberculatus, M. flagellaris, N. orbicularis, Phaeoceros* sp*., Phaeomegaceros chiloensis* and *Phymatoceros phymatodes*, together covering all the hornwort families. It is important to note that these hornwort species are deeply diverged; they share the most recent common ancestor almost 300 million years ago (Bechteler et al., 2023), which predates the origin of flowering plants (Ramírez-Barahona et al., 2020). The successful transient transformation of multiple hornwort species suggests that our method is widely applicable across taxa, thus broadening the scope of genetic manipulation in hornworts. The ability to transform diverse hornwort species enables comparative studies to identify conserved genetic elements, especially for convergently evolved traits. For example, pyrenoids have originated independently approximately five to six times during hornwort evolution, resulting in vastly different morphologies and CCM strength across species (Vaughn et al., 1990; Hanson et al., 2002; Villarreal & Renner, 2012; Li et al., 2017). Understanding the genetic basis of such repeated evolution could have important implications in engineering CCMs in other plants. Likewise, U/V sex chromosomes have experienced multiple gains and losses in hornworts (Villarreal & Renner, 2013). Future work could use our biolistics method to test if similar genetic elements have been convergently recruited to sex chromosomes.

Despite the relatively high number of transient events, we had mixed success in generating stable transformants from diverse hornwort species. While 29 *A. fusiformis* stable lines were recovered from one experiment, we did not recover stable events from the remaining species. It is likely that the method will need to be optimized for each species. In particular, we have observed that a number of species have a slower growth rate on AG medium, which indicates that the post homogenization recovery time may need to be extended. In our constructs, the *A. agrestis Ef1α* promoter drives *hpt* expression, however, it is possible that this promoter may not function well in other hornwort species, especially considering the timing of divergence (Fig. 4), and their native *Ef1a* promoter is required. The optimal antibiotic concentration to select for transgenic events may also vary between species.

### Relative high frequency of low transgene copy number in stably transformed lines

A common issue for biolistics-mediated transformation is the possibility of multiple transgene insertions into the genome. Here we report that 33% of sequenced transgenic lines were estimated to have single-copy insertions, and 50% of the lines have less than four copies. The frequency of low-copy insertions is much higher when compared to biolistics transformation of other bryophytes. Studies in the liverwort *Marchantia polymorpha* and the moss *Physcomitrium patens* found that only 10% of regenerated lines had single copies of the transgene (Irifune et al., 1996; Šmídková et al., 2010). Nevertheless, we did occasionally observe lines with a large transgene copy number, with two lines having an estimated 63 and 73 copies, respectively. While *Agrobacterium*-mediated transformation of hornworts has been demonstrated, the frequency of low copy number events was not reported (Frangedakis et al., 2021b, Waller et al., 2023). It is likely that the frequency of single-copy insertions can be further increased using a minimal cassette, where the genes of interest are amplified using PCR, eliminating the backbone from transformation. This technique typically requires less DNA for transformation. Two studies in wheat aimed to increase the number of single copy events by using minimal cassettes, resulting in an average of 38% and 50% of single-copy insertions (Tassy et al., 2014; Ismagul et al., 2018), while in rice 80% of regenerated lines were estimated to have less than two transgene copies (Fu et al., 2000).

### Using fluorescent protein tagging to visualize hornwort pyrenoids

Application of this method enabled us to observe localization of fluorescently tagged proteins within 48 hours post bombardment and nine days from the point the tissue was first homogenized. The biolistics method thus provides the ability to quickly assess protein function, localization, and interactions without the need to generate stable transformants. We demonstrated that our method can be used to study proteins associated with pyrenoids and CCM function.

Pyrenoids are membraneless micro-compartments within the chloroplasts of algae and hornworts that are composed predominantly of Rubisco. This aggregation of Rubisco enables the function of a biophysical CCM providing a central locus in which to concentrate CO_2_. While pyrenoids are present in many algal species and have been extensively studied in the model alga *Chlamydomonas reinhardtii* (He et al., 2023), hornworts are the only land plants with pyrenoids and have received little research attention (Li et al., 2017). Early ultrastructural studies reported that hornwort pyrenoids are tightly sandwiched by thylakoids (as opposed to starch sheaths in algae) (Burr, 1970; Vaughn et al., 1992), and based on immunogold labeling are indeed the locations where Rubisco are concentrated (Vaughn et al., 1990). We set out to visualize pyrenoids by fluorescent protein tagging and to establish pyrenoid marker proteins for future studies. To this end, we focused on the Rubisco small subunit (RbcS) as well as Rubisco activase (RCA), which is required for Rubisco activation and activity maintenance (McKay et al., 1991). In *C. reinhardtii* both proteins are known to localize predominantly in pyrenoids (Mackinder et al., 2017). We co-transformed RbcS and RCA tagged with the mScarlet-I and mVenus fluorescent protein, respectively, into *A. agrestis* thalli. RbcS and RCA were found to clearly co-localize to pyrenoids based on the overlapping mScarlet-I and mVenus signals (Fig. 5). To our knowledge, this is the first application of genetic engineering to interrogate gene function in hornworts. Importantly, it is now possible to use either fluorescently tagged RbcS or RCA as a pyrenoid marker, and co-transform it with candidate genes to build a spatial protein localization model for hornwort CCMs. Furthermore, we show the successful regeneration of lines stably expressing either *RbcS:mScarlet-I* or *RCA:mVenus*. Currently, these transformants are being used to track pyrenoid development and to understand how it is impacted by environmental factors such as CO_2_ concentration.

In conclusion, we report the first biolistics protocol for transformation of a phylodiverse set of hornwort species, as well as the first gene functional study in the model hornwort *Anthoceros agrestis*. We anticipate our work to allow new avenues for investigation and biotechnological applications, such as genome editing. The achievement of both stable and transient transformation provides powerful tools for studying gene function in hornworts. These advances lay the foundation for future research aimed at unraveling the mysteries of hornwort biology and harnessing their unique characteristics for sustainable agriculture.

## Supplemental Data

Supplemental Table 1. BUSCO genes used in transgene copy number estimation analysis.

Supplemental Figure 1. Correlation between manual and CellProfiler counting of cells transiently expressing GFP.

Supplemental Figure 2. *Anthoceros agrestis* stable lines regenerated from experiment 1.

Supplemental Figure 3. *Anthoceros agrestis* stable lines regenerated from experiment 2.

Supplemental Figure 4. *Anthoceros agrestis* stable lines regenerated from experiment 3.

Supplemental Figure 5. *Anthoceros fusiformis* stable lines.

## Abbreviations

BUSCO: Benchmarking Universal Single-Copy Orthologue
CCM: CO_2_-concentrating mechanism
Ef1α: Elongation factor 1 alpha
GFP: green fluorescent protein
HPT: hygromycin phosphorylase
RbcS: Rubisco small subunit
RCA: Rubisco activase
Rubisco: ribulose-1,5-bisphosphate carboxylase/oxygenase

## Acknowledgements

We thank Eftychis Frangedakis for providing the materials in the OpenPlant toolkit and transformation advice, Peter Schafran for helping with estimating transgene copy numbers, Mamta Srivastava for helping with confocal and fluorescence microscopy and Jenna Sins and Evan Smith for laboratory assistance.

## Author contributions

FWL and JVE: conceptualization; DL: formal analysis; FWL, JVE, and LHG: funding acquisition; DL, TAR, and AG: investigation; DL and AG: methodology; FWL, JVE, and LHG: supervision; DL and TAR: visualization; DL and FWL: writing—original draft.

## Data availability

The raw Illumina reads for quantifying transgene copy numbers were deposited in NCBI SRA (under the BioProject PRJNA1029596).

